# Estimating Heritability and Genetic Correlation in Case Control Studies Directly and with Summary Statistics

**DOI:** 10.1101/256388

**Authors:** Omer Weissbrod, Jonathan Flint, Saharon Rosset

## Abstract

Methods that estimate heritability and genetic correlations from genome-wide association studies have proven to be powerful tools for investigating the genetic architecture of common diseases and exposing unexpected relationships between disorders. Many relevant studies employ a case-control design, yet most methods are primarily geared towards analyzing quantitative traits. Here we investigate the validity of three common methods for estimating genetic heritability and genetic correlation. We find that the Phenotype-Correlation-Genotype-Correlation (PCGC) approach is the only method that can estimate both quantities accurately in the presence of important non-genetic risk factors, such as age and sex. We extend PCGC to work with summary statistics that take the case-control sampling into account, and demonstrate that our new method, PCGC-s, accurately estimates both heritability and genetic correlations and can be applied to large data sets without requiring individual-level genotypic or phenotypic information. Finally, we use PCGC-S to estimate the genetic correlation between schizophrenia and bipolar disorder, and demonstrate that previous estimates are biased due to incorrect handling of sex as a strong risk factor. PCGC-s is available at https://github.com/omerwe/PCGCs.

## Introduction

Much of the theory underlying methods for estimating two key measures of disease genetic architecture, heritability and genetic correlation, was designed for cohort studies of quantitative phenotypes. Consequently, when applied to studies of categorical traits, these methods may contain unacknowledged biases that may affect the accuracy of the estimates.

The problem of accurately estimating heritability and genetic correlation is usually translated into questions about variance and covariance components in properly defined mathematical models. A commonly held misconception states that variance components can be accurately calculated in case-control studies by virtue of applying a correction factor to results derived under a quantitative trait framework (e.g. refs.^1–4^). However, this is not true when risk factors (including risk variants) exert a strong influence on disease risk. In this paper we examine the validity of approaches for estimating heritability, covariance and correlation (covariance standardized to a [-1, 1] scale) in case-control studies of disease.

Broadly speaking, there are three common approaches for carrying out these tasks. The first is based on restricted maximum likelihood estimation (REML) in the linear mixed model (LMM)^5^ framework, and is implemented in some widely used tools^6,7^. This approach has been extensively applied to heritability estimation^2,7,8^ and more recently to genetic correlation estimation^7,9–11^.

The second approach is based on regression of phenotype correlations on genotype correlations and relies on less restrictive assumptions than the LMM approach. It is broadly known as Haseman-Elston (HE) regression^12,13^. For estimating heritability in case-control studies of disease, the relevant variant is called PCGC^14,15^. In this paper we extend PCGC to also estimate genetic correlation.

The third approach we consider is the family of linkage disequilibrium score regression (LDSC) methods, which estimate heritability and correlation while accounting for LD^1,16^, and have recently been applied to several large scale studies of psychiatric disorders^17,18^. LDSC is attractive because it requires only publicly available summary statistics from genetic studies, thereby avoiding privacy and logistical concerns^19^. Although other summary-statistics based methods have been proposed recently, we focus on LDSC, as alternative methods cannot be applied in the presence of LD^3^ or are not directly designed for analysis of categorical phenotypes^4,20,21^.

Here we examine all three approaches and demonstrate that LDSC and REML can yield biased estimates in the presence of covariates representing major risk factors such as sex and age, while PCGC remains accurate. We further develop a new version of PCGC that can work with summary statistics that explicitly take the case-control sampling into account. We demonstrate the value of this method by investigating the genetic correlation between schizophrenia and bipolar disorder, and between type 1 diabetes and coronary artery disease. We demonstrate that the estimates of both quantities are severely biased under alternative methods due to incorrect handling of sex – an important risk factor. Finally, we provide best practice recommendations depending on the available data and the trait characteristics.

## Materials and Methods

### Underlying Mixed Effects Model

We adopt the theoretical framework of the liability threshold model^22,23^, which assumes that every individual *i* has a normally distributed latent liability value for trait *t*, 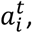 such that cases of trait *t* are individuals whose liability exceeds a given cutoff.

We additionally assume that the liability of trait *t* can be decomposed into a genetic and an environmental term,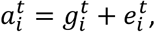 such that 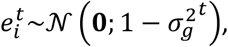, and the vector of genetic effects in a study, 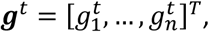, follows a multivariate normal distribution:

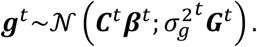

Here, ***C***^*t*^ is a design matrix of covariates, ***β***^*t*^ is a column vector of fixed effects,***G***^*t*^ is a matrix of kinship coefficients, and 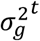 is a genetic variance parameter. Under these assumption, every individual *i* has an observed binary affection status indicator for trait *t*, 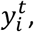 such that 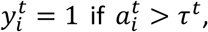 where 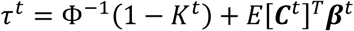is the affection cutoff for trait *t* with prevalence *K*^*t*^ and where Φ,^−1^(·) is the inverse cumulative standard normal density.

For a pair of traits *t*_1_, *t*_2_, the concatenated liabilities vector follows a multivariate normal distribution,

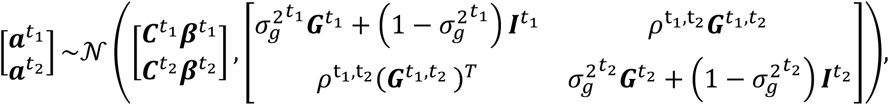

where ***G***^*t*_1_, *t*_2_^ is the matrix of between-study kinship coefficients, *ρ*^*t*_1_, *t*_2_^ is the genetic covariance, and ***I***^*t*_1_^, ***I***^*t*_2_^ are identity matrices.

The quantities we investigate in this paper are defined as follows:

a. The heritability of trait *t*, defined as 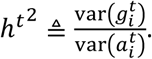
b. The genetic covariance of two traits *t*_1_, *t*_2_, defined as 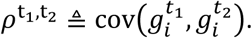
c. The genetic correlation of two traits *t*_1_, *t*_2_, defined as 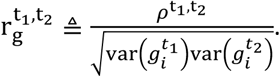

### The Effect of Ignoring Covariates

The main contribution of PCGC-s over LDSC is its ability to account for covariates. Although it is rarely possible to measure all possible covariates affecting the trait of interest, covariates with a strong effect (such as the effect of sex on coronary artery disease) are often measured. This raises the question whether omission of such important covariates affects heritability and genetic correlation estimates. We provide a proof in the Supplementary Material that omission of covariates does not bias these estimates if the covariate effects are normally distributed and are uncorrelated with the genetic effect. The main idea behind the derivation is that the environmental effect represents the aggregated effect of unmeasured covariates, and can thus absorb the effect of omitted covariates when these assumptions hold.

The assumption of normality approximately holds if a trait is influenced by a large number of covariates with small effects, owing to the central limit theorem. However, many traits are strongly influenced by a small number of non-normally distributed covariates, such as sex. Heritability estimates with omitted covariates can become inaccurate in the presence of such strong covariates. In contrast, genetic correlation is accurately estimated in the simulations even in the presence of strong non-normal covariates, suggesting that the errors in the estimation of genetic covariance and genetic variance approximately cancel out when dividing one by the other. However, this observation is currently unsupported by statistical theory.

The assumption that covariates are uncorrelated with the genetic effect is often violated when using heritable covariates, such as genetic principal components. This problem can easily be circumvented by regressing the omitted covariates out of the genotypes and correcting the individual-level affection cutoffs prior to parameter estimation or to computing summary statistics (Supplementary Material). We caution that regression of covariates out of binary phenotypes as suggested in ref.^24^ can yield incorrect estimates in case control studies, even for the genetic correlation (as verified in the results section).

### Marginal and Conditional Heritability

An important point often overlooked in heritability estimation is that covariates such as sex and age also contribute to the liability variance. Since the liability is non-identifiable, it is typically assumed to have a unit variance when conditioning on measured covariates. In this case, the marginal liability variance is given by 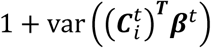 and consequently, heritability is given by 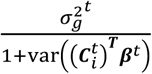 (Supplementary Material). Alternatively, one could assume that the marginal variance is one, in which case the conditional variance is smaller than one.

In contrast, many studies define the genetic variance 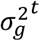 as the heritability, even in the presence of covariates. We therefore denote the former definition as marginal heritability and the latter definition as conditional heritability, because the latter definition uses the variance of the liability conditional on measured covariates.

In this paper we consider marginal heritability, both because this definition is arguably more natural as different studies using different covariates are ultimately interested in estimating the same quantity, and because we show via simulations (Supplementary Figure S1) that LDSC tends to severely underestimate the conditional heritability (as compared to less severe overestimation of marginal heritability -Figure 1). Therefore, we do not consider estimation of conditional heritability further in this paper.

**Fig. 1:**
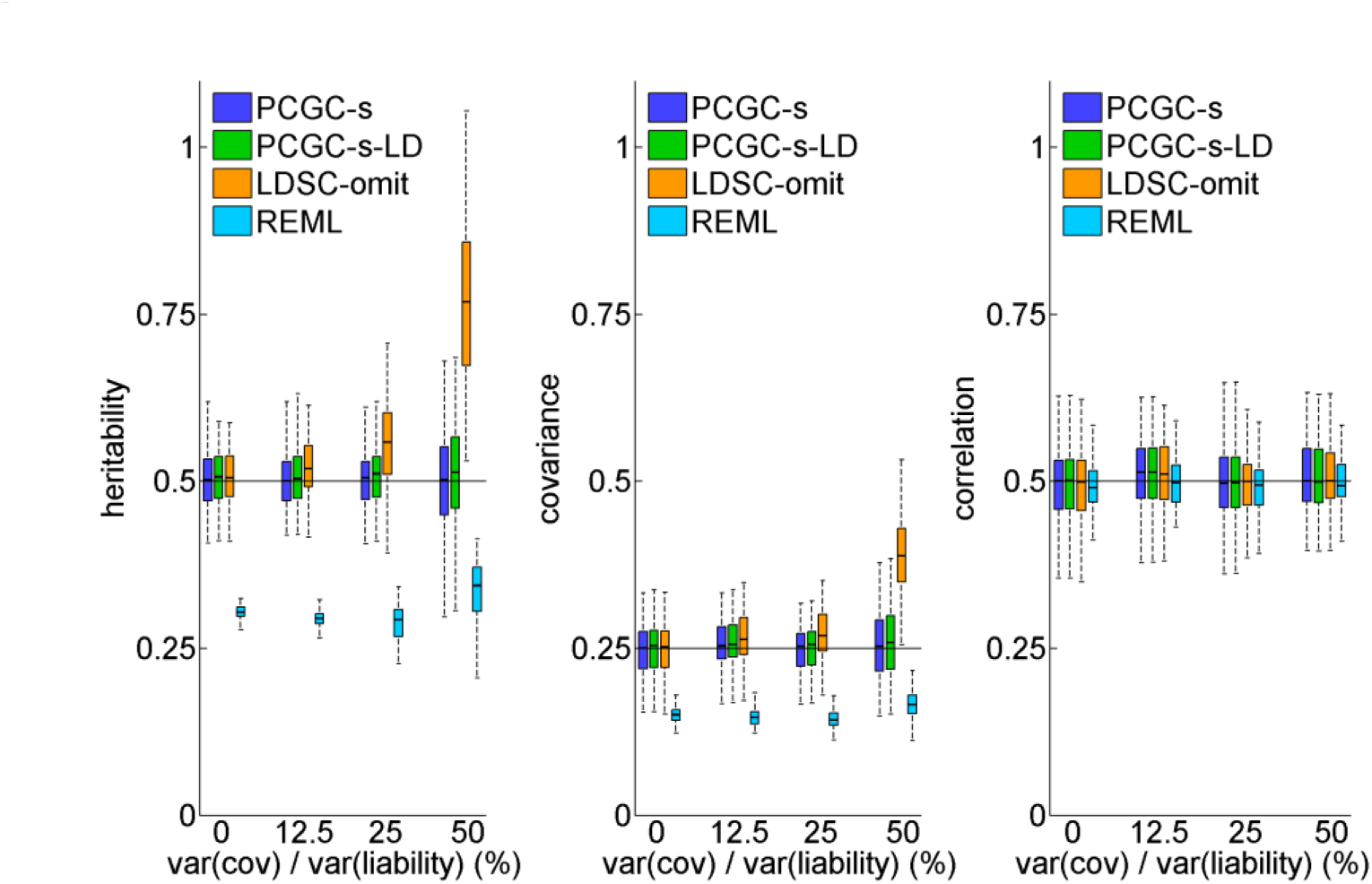
The effect of covariate strength. PCGC-s and PCGC-s-LD estimate all parameters accurately under all settings; LDSC-omit estimates of heritability and genetic covariance become increasingly inaccurate as the covariates strength increases; REML misestimates heritability and genetic covariance under all settings. All methods estimate genetic correlation accurately. The black horizontal lines indicate the true parameter values. 100 experiments were performed for each unique combination of settings, and each study included 2,000 cases and 2,000 controls.

### PCGC-s with no Covariates

PCGC with no covariates estimates *ρ*^*t*_1_, *t*_2_^ by regressing standardized phenotypic correlations 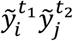 on kinship coefficients 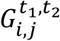 and then dividing the resulting estimator by a constant *ƒ*(*t*_1_, *t*_2_) (Supplementary Material). This estimation encapsulates both genetic covariance and heritability, which for a trait *t* with no covariates is given by *ρ*^*t*, *t*^.

The PCGC estimator can be computed without individual-level data by using the following two summary statistics:

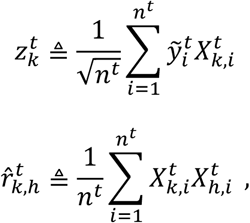

where *n*^*t*^is the sample size of study *t* and 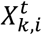 is the value of the *k*^th^ variant of individual *i* in study *t*, after standardization. It is also possible to use logistic regression-based or other types of summary statistics, but this constitutes an approximation (Supplementary Material).

Using these quantities and denoting *S*^*t*_1_, *t*_2_^ as the set of all pairs of indices *i, j* that refer to the same individual shared between the two studies, the PCGC estimator can be written as:

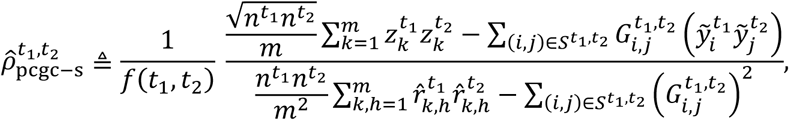

where *m* is the number of variants, and *ƒ*(*t*_1_, *t*_2_) is given by:

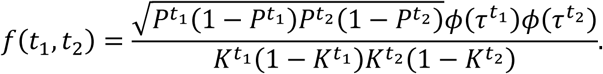

Here, *k*^*t*^ and *p*^*t*^ are the prevalence of trait *t* and the case-control proportion of study *t*, respectively,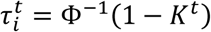 is the liability cutoff, and *ϕ*(·), Φ(·) are the density and cumulative distribution of the standard normal distribution, respectively.

The resulting estimator approximately coincides with the LDSC estimator if there are no overlapping individuals and the in-sample LD estimates in both studies are the same as in the reference population used by LDSC^24^. The extension to estimating multiple variance components is straightforward (Supplementary Material).

The second term of the numerator and of the denominator can be computed by research groups with access to the genotypes and phenotypes of overlapping individuals, which often consist of control cohorts, or can be approximated via the approximation 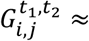 1.0 for overlapping individuals, as done implicitly in LDSC^2^. However, we caution that even minor deviations (which can occur for example by regressing out principal components) can affect the approximation (Supplementary Material).

A particularly convenient property of 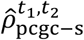 in the absence of covariates is that when estimating the genetic correlation, all terms dependent on the trait prevalence vanish. This is convenient because the true trait prevalence is often not known with certainty.

### PCGC-s with Covariates

In the presence of covariates, PCGC estimates *ρ*^*t*_1_, *t*_2_^ by regressing 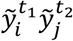 on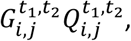 where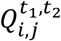 is a quantity that depends on the covariates of individuals *i* and *j*, and so the regression constant is different for every pair of individuals^14^. The corresponding PCGC-s estimator is given by:

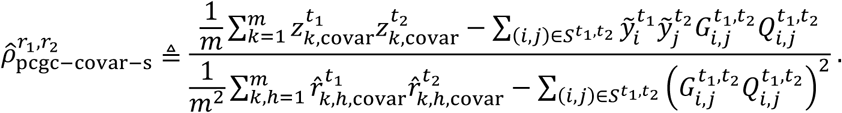

The above quantities are defined as follows:

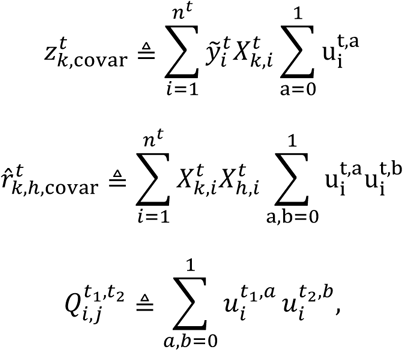

where 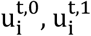 are given by:

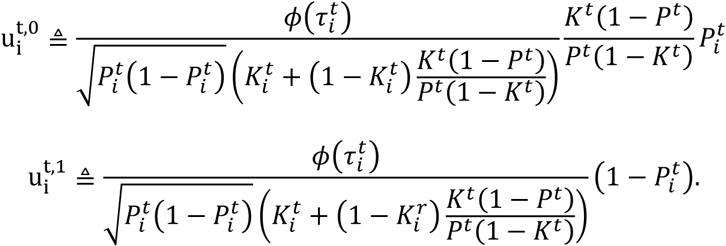

Here,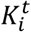is the probability of individual *i* being a case conditional on her covariates, 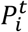 is the probability of individual *i* being a case conditional on her covariates and on being ascertained into the study, and 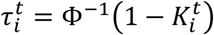 is the liability cutoff of individual *i* conditional on her covariates.

The full derivation, an extension for multiple variance components and an approximation that requires a single summary statistic instead of using 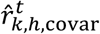 (which requires a number of statistics equal to the number of pairs of variants) are provided in the Supplementary Material.

As in the case of no covariates, the second term of the numerator and denominator can be computed by research groups with access to overlapping individuals, which often consist of control cohorts. Third parties with no access to overlapping individuals can approximate the terms on the right-hand sides of the numerator and the denominator given appropriate summary statistics (Supplementary Material).

## Results

We assume that quantitative binary traits are governed by the liability threshold model, which postulates that every individual is associated with a latent normally distributed variable called the liability, such that individuals with liability greater than a given cutoff are cases and the rest are controls. The common practice is to define heritability and genetic covariance on the liability scale rather than the observed scale. We are therefore interested in estimating the following quantities (see Methods for exact definitions): (a) Heritability the fraction of liability variance explained by genetics; (b) Genetic covariance - the covariance between the genetic components of two traits on the liability scale, and (c) genetic correlation - the genetic covariance standardized to a [-1,1] scale.

We are concerned with the three following questions:

1. Can quantities (a)-(c) be estimated reliably given genotypic and phenotypic data?
2. Can quantities (a)-(c) be estimated reliably given summary statistics via LDSC?
3. Can quantities (a)-(c) be estimated reliably given summary statistics via an alternative method?

The answers to questions 1,2 are summarized in Table 1. Briefly, PCGC is the only method that can estimate all quantities of interest under all investigated settings. REML provides inconsistent estimates of quantities (a)-(b), and empirically provides consistent estimates of quantity (c). LDSC can provide consistent estimates of quantities (a)-(b) in the absence of covariates, and provides consistent estimates of quantity (c) when no covariates are included in the analysis. To answer question 3, we present a reformulation of PCGC called PCGC-s that can estimate quantities (a)-(c) reliably using only summary statistics, both with and without covariates (Methods).

**Table 1:**
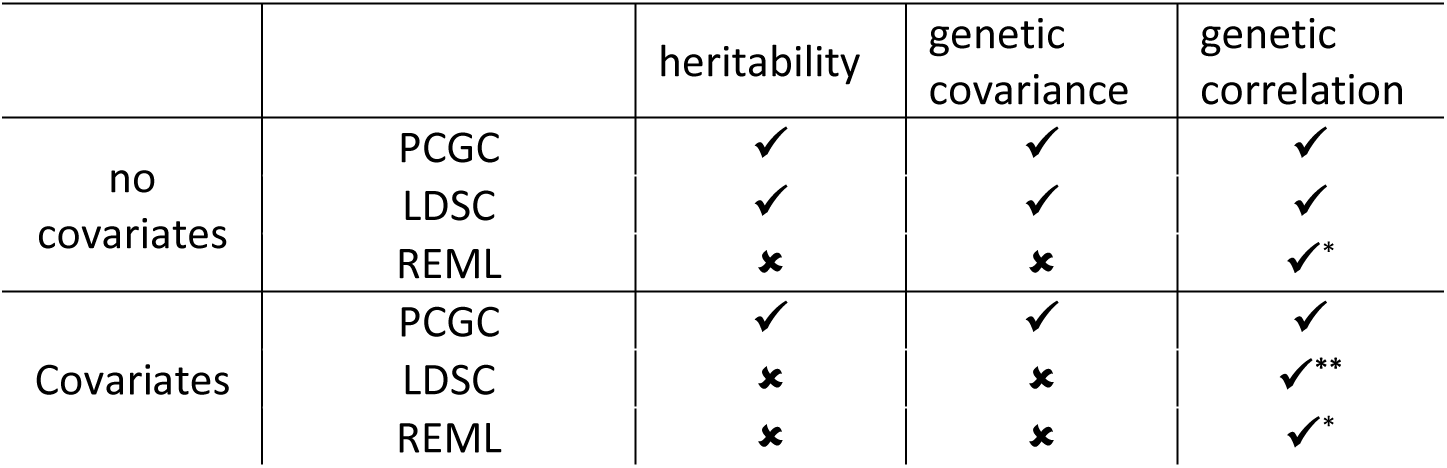
Estimation correctness of the investigated methods. LDSC behaves differently depending on whether covariates are present. The entries marked with * indicate that although the estimated quantity is empirically unbiased in simulations, it is given by the division of two biased estimates (the estimate in the second column divided by the square root of the estimate in the first column), suggesting that estimation errors cancel each other in the division. We are not currently aware of a theoretical justification for this behavior. The entry marked with ** is only empirically correct as long as covariates are excluded from the analysis.

|  |  | heritability | genetic covariance | genetic correlation |
| --- | --- | --- | --- | --- |
| no covariates | PCGC | ✓ | ✓ | ✓ |
|  | LDSC | ✓ | ✓ | ✓ |
|  | REML | ✗ | ✗ | ✓* |
| Covariates | PCGC | ✓ | ✓ | ✓ |
|  | LDSC | ✗ | ✗ | ✓** |
|  | REML | ✗ | ✗ | ✓* |

### Simulation Studies

We conducted simulation studies to investigate the behavior of the evaluated methods in case-control studies; such simulations require first obtaining a very large pool with hundreds of thousands of individuals, and then sampling a small fraction of cases according to the trait prevalence^2,14,16,25^. ded in Supplementary Material.

Our simulations span a wide range of scenarios, with various levels of prevalence, heritability, genetic correlation, sample sizes, number of single nucleotide polymorphisms (SNPs), number of covariates, LD patterns, fraction of shared controls, and trait polygenicity. In each experiment we varied one or more of the above parameters while keeping the others fixed. The default simulation parameters used 1% prevalence, 50% heritability and 50% genetic correlation, with each study having 2,000 cases, 1,000 unique and 1,000 overlapping controls, and 10,000 SNPs with a correlation of approximately 95% between adjacent SNPs. In most simulations all SNPs influenced the phenotype, though we verified that relaxing this assumption does not affect the results (see details below). 100 simulations were conducted for each unique combination of settings.

The examined methods included (a) PCGC-s; (b) PCGC-s-LD, which is an approximate version of PCGC-s that uses external LD estimates (but uses data about overlapping individuals; Supplementary Material); (c) LDSC with omitted covariates (LDSC-omit), and (D) REML, using the implementation in GCTA^6^ (exact execution details are provided in the Supplementary material). Note that PCGC-s is exactly equivalent to PCGC when all required summary statistics are provided. LDSC-omit refers to LDSC that does not include any covariates in the analysis, and was used because explicit inclusion of covariates can lead to highly biased estimates, as demonstrated below. In most simulations, LDSC-omit was based on our own implementation, to avoid confounding the analysis by implementation details. Specifically, our implementation of LDSC-omit used a predetermined intercept and did not weight summary statistics by their posterior variance, similarly to PCGC-s-LD (see Discussion for elaboration on these issues). In additional simulations described below, we demonstrated that when using the ldsc software instead of our own implementation, LDSC-omit became less accurate.

Our first experiment examined the impact of covariate effect magnitude on the estimation of heritability, genetic covariance and genetic correlation. We simulated data sets with 5 binary covariates that explained various fractions of the liability variance, where the first covariate accounted for 95% of the aggregated covariates effect. All methods estimated correlation well, but PCGC-s and PCGC-s-LD were the only methods that estimated the two other quantities accurately (Figure 1). Both PCGC-s and PCGC-s-LD had a statistically significant advantage over LDSC-omit in heritability estimation (P<2.10 × 10^−2^, P<1.67 × 10^−6^, P<6.45 × 10^−24^ for covariates that explained 12.5%, 25% and 50% of the liability variance, respectively; Binomial test for PCGC-s-LD; PCGC-s results were effectively the same). The accuracy of LDSC-omit improved as effect sizes became smaller; LDSC-omit and PCGC give very similar estimates in the absence of covariates, as expected from theory (Supplementary Material). REML consistently underestimated heritability despite using the correction for case-control ascertainment implemented in GCTA^2^. We note that the extent of under-estimation by REML is not fixed with a known ratio but depends on various parameters^14^. We also obtained similar results when ignoring the contribution of covariates to the liability variance (Methods, Supplementary Figure S1).

The next experiment examined the implications of having normal versus non-normal covariate effects, by considering three settings: (a) A single binary covariate; (b) a single normally distributed covariate, and (c) 20 equally strong binary covariates. In all settings the covariates jointly explained 40% of the liability variance. Setting (a) encodes a non-normal aggregated effect, whereas settings (b) and (c) encode a normal and an approximately normal effect (owing to the central limit theorem), respectively. In setting (a) LDSC-omit was substantially less accurate than PCGC-s (P<3.21 × 10^−19^; Binomial test) and PCGC-s-LD (P<2.73 × 10^−20^; Binomial test), because its underlying model is violated in the presence of strong non-normally distributed covariates (Figure 2, Methods). The bias of LDSC-omit decreased when decreasing the magnitude of the covariate effects, similarly to the results shown in Figure 1.

**Fig. 2:**
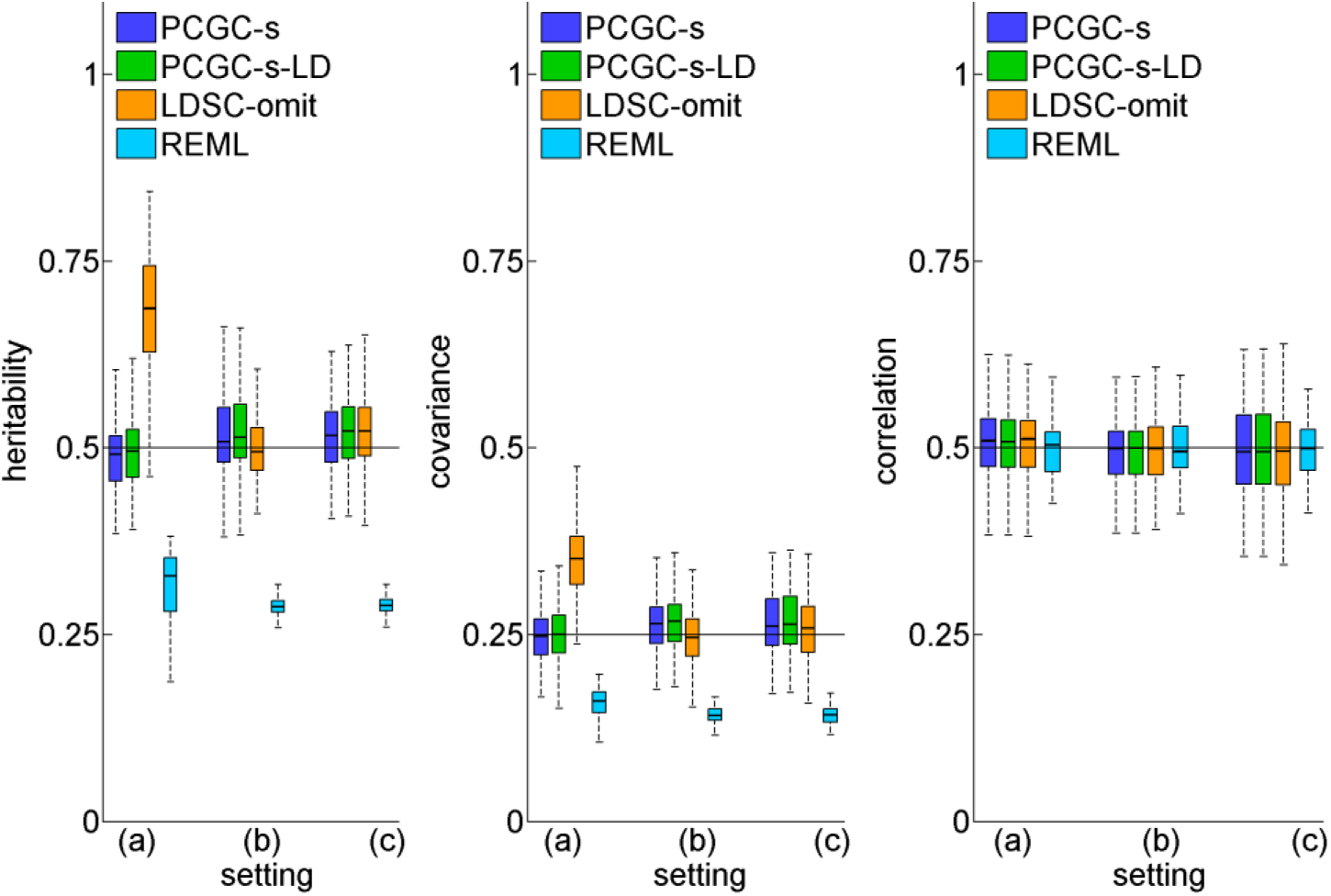
The effect of the covariate effects distribution. Setting (a) includes a single binary covariate; Setting (b) includes a single normally distributed covariate; Setting (c) includes 20 binary variables with equal strength, yielding an approximately normal aggregated effect owing to the central limit theorem. PCGC-s and PCGC-s-LD are the only methods that accurately estimate heritability and genetic covariance in setting (a), where the covariates effects distribution is far from normal. 100 experiments were performed for each unique combination of settings, and each study included 2,000 cases and 2,000 controls.

In additional experiments, we simulated data with one strong and four weak binary covariates as in the first experiment, where the covariates jointly explained 25% of the liability variance, and verified that the results remained similar under various levels of heritability (Supplementary Figure S2), genetic correlation (Supplementary Figure S3), prevalence (Supplementary Figure S4), LD (Supplementary Figure S5), fraction of shared controls (Supplementary Figure S6), numbers of covariates (Supplementary Figure S7), sample sizes (Supplementary Figure S8), numbers of simulated causal SNPs (Supplementary Figure S9), and trait polygenicity (Supplementary Figure S10). We also explored running LDSC-omit using the ldsc software (Supplementary Figure S11), and using logistic regression-based summary statistics (Supplementary Figures S12-S13).

We also examined the effect of using LDSC without omitting covariates, by regressing measured covariates out of the phenotypes and genotypes prior to computing summary statistics, as recommended in refs.^16,24^ Our results demonstrate that LDSC estimates are severely down-biased in this setting, with an average bias of over 10% in heritability and covariance estimation, and of over 5% in correlation estimation, under realistic settings (Supplementary Figures S14-S15).

Finally, we verified that PCGC-s-LD is highly computationally efficient. Since PCGC-s-LD uses only summary statistics, it can perform estimation for data with millions of variants and hundreds of thousands of individuals in less than five minutes (results not shown).

### Estimating the genetic architecture of schizophrenia and bipolar disorder

To demonstrate the behavior of the methods on real data we studied the heritability and genetic correlation of schizophrenia (SCZ)^26^ and bipolar disorder (BP)^27^. To prevent confounding due to population stratification^28^, we restricted the analysis to two highly concordant Swedish data sets consisting of 1,745 SCZ cases, 1,268 BP cases and 6,293 controls, 2,566 of which are shared between the studies^26,27^ (Supplementary Material). The covariates included 10 principal components and sex, which is a major risk factor for both diseases.

The PCGC-s heritability estimates for SCZ and BP were 39.2% and 41.7%, respectively. The estimated genetic correlation was 42.4%, which is substantially lower than previous estimates of 68% using REML^10^, and 79% using LDSC^1^. We further verified that when omitting the covariates, the PCGC-s estimates increased to 60%, suggesting that incorrect treatment of non-genetic risk factors can lead to inflated estimates. When invoking LDSC on the same data using the ldsc software, the estimated correlation could not be computed when omitting covariates due to negative estimated heritabilities, and was 16.9% when regressing the covariates out of the phenotypes (Table 2). We conclude that improper handling of covariates and of sample overlap in case-control studies can lead to substantially biased estimates and to incorrect conclusions regarding the genetic architecture of genetic diseases.

**Table 2:**
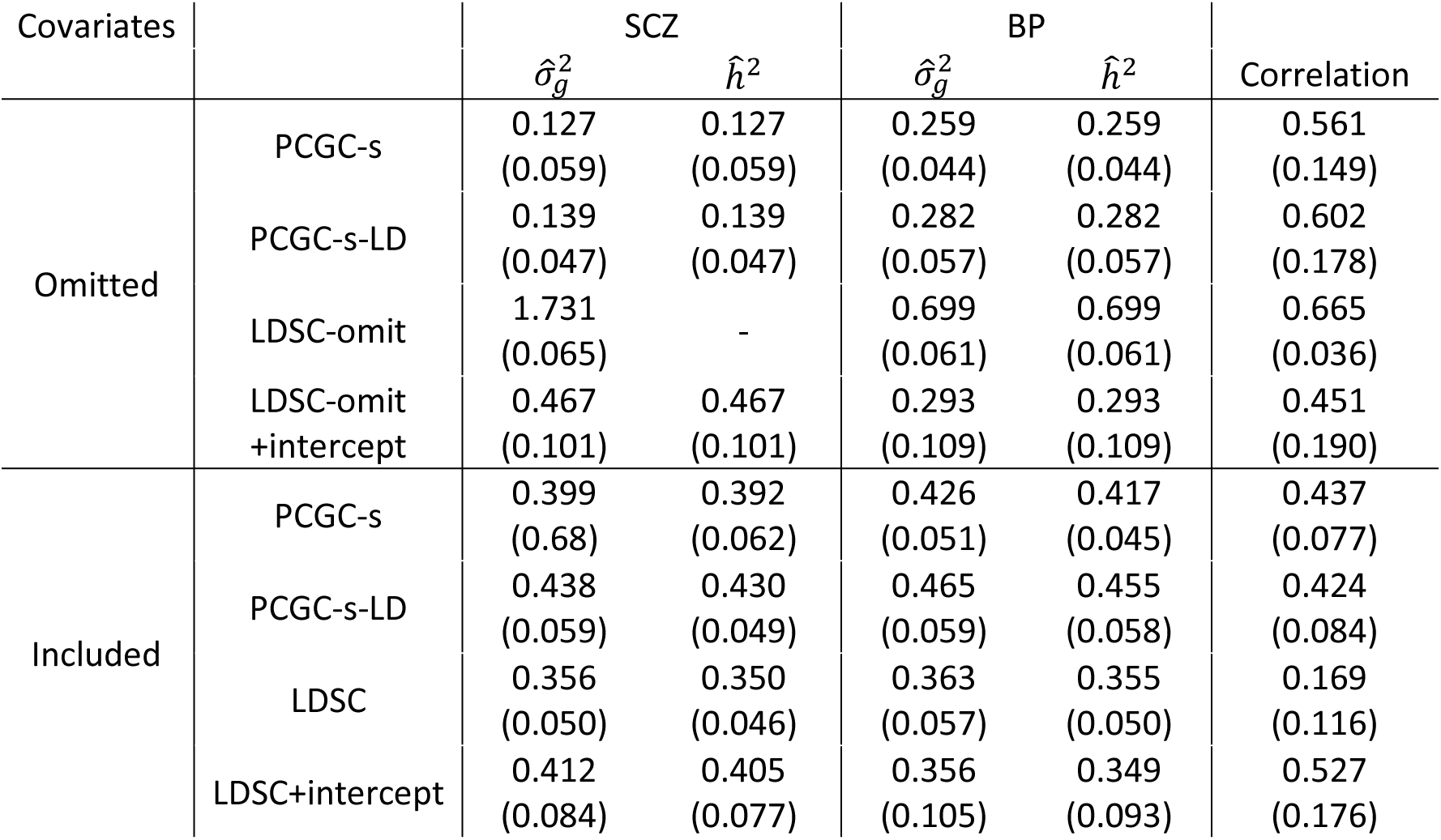
Results of real data analysis of psychiatric disorders. Shown are the estimated values of the genetic variance 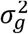 (also termed the conditional heritability in this paper), the marginal heritability *h*^2^(which is equal to 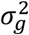when no covariates are present, and smaller than 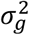 in the presence of covariates) and the genetic correlation. Standard errors were computed via a block jackknife of 200 blocks of consecutive SNPs, as in LDSC. LDSC+intercept is the LDSC estimator when fitting an intercept from the data^2^. LDSC-omit is different from PCGC-s-LD with omitted covariates because of differences in the predetermined intercept value due to normalization (Supplementary Material). LDSC results were computed using the ldsc software. Values marked with “-” could not be computed because of negative or illegal parameter estimates.

### Estimating the genetic architecture of type 1 diabetes and coronary artery disease

To further evaluate PCGC-S, we studied the correlation between type 1 diabetes (T1D) and coronary artery disease (CAD), using data from the Wellcome trust case control consortium 1 (WTCCC1)^29^. It is known that T1D is associated with an increased risk for CAD^30^, but the role of genetics in this association is not clear. We chose to explore this example because of the expected impact of covariates on the result: T1D is very strongly affected by SNPs in the major histocompatibility complex (MHC) region, and sex is a major risk factor for CAD. We thus modeled the effects of these risk factors as fixed rather than random, and investigated the implications of inclusion and exclusion of these covariates. The analysis details are provided in the Supplementary Material.

The results demonstrated the existence of a positive genetic correlation between T1D and CAD, and corroborated the simulation studies (Table 3, Supplementary Table 1). As expected, inclusion of covariates had a minor effect on PCGC-s estimates, decreasing the heritability estimate for T1D from 23.7% to 18.3%, and for CAD from 40.5% to 39.9%, and slightly increasing the genetic correlation estimate from 18.1% to 19.2%. The LDSC heritability estimates for T1D and CAD when omitting covariates (35% and 58.8%, respectively) were greater than those of PCGC-s (consistent with our simulation results) and the correlation estimate was also greater (28.4%). LDSC heritability estimates were nonsensical (non-positive or greater than one) when including covariates or fitting an intercept rather than using a predetermined one. REML estimation of genetic correlation using gcta failed to converge.

**Table 3:** Results of real data analysis of T1D and CAD. The table fields are the same as in Table 2. LDSC results are based on our own implementation to provide a detailed comparison with PCGC-s that is not confounded by implementation details. Results using the ldsc software are provided in Supplementary Table 1.

| Covariates |  | T1D |  | CAD |  | Correlation |
| --- | --- | --- | --- | --- | --- | --- |
| | | $\hat{\sigma}_g^2$ | $\hat{h}^2$ | $\hat{\sigma}_g^2$ | $\hat{h}^2$ | |
| Omitted | PCGC-s | 0.237<br>(0.044) | 0.237<br>(0.044) | 0.405<br>(0.063) | 0.405<br>(0.063) | 0.181<br>(0.115) |
|  | PCGC-s-LD | 0.245<br>(0.045) | 0.245<br>(0.045) | 0.420<br>(0.065) | 0.420<br>(0.065) | 0.181<br>(0.115) |
|  | LDSC-omit | 0.350<br>(0.046) | 0.350<br>(0.046) | 0.588<br>(0.066) | 0.588<br>(0.066) | 0.284<br>(0.074) |
|  | LDSC-omit<br>+intercept | 0.013<br>(0.105) | 0.013<br>(0.105) | 0.020<br>(0.109) | 0.020<br>(0.109) | - |
| Included | PCGC-s | 0.241<br>(0.066) | 0.183<br>(0.050) | 0.435<br>(0.070) | 0.399<br>(0.062) | 0.192<br>(0.139) |
|  | PCGC-s-LD | 0.250<br>(0.069) | 0.190<br>(0.052) | 0.451<br>(0.065) | 0.413<br>(0.060) | 0.191<br>(0.139) |
|  | LDSC | -1.75<br>(0.038) | - | -0.33<br>(0.058) | - | - |
|  | LDSC+intercept | -0.03<br>(0.046) | - | -0.07<br>(0.09) | - | - |

We conclude that accounting for covariates can substantially affect heritability and genetic correlation estimates. However, we caution that the results are sensitive to preprocessing of the data (Supplementary Tables S2-S4, Supplementary Material; see Discussion). We also present genetic correlation estimates between all phenotypes included in the WTCCC1 study, confirming some well known significant correlations, such as between hypertension and coronary artery disease; and others that have been tentatively suggested in the literature, such as between rheumatoid arthritis and coronary artery disease^31,32^(Supplementary Table S5).

## Discussion

Our major conclusions regarding the existing approaches can be summarized as follows: (i) REML severely misestimates heritability and genetic covariance in case-control studies under all settings (as has been pointed out previously^7,14,25^). In settings without binary covariates REML accurately estimates genetic correlation, but it becomes biased in the presence of such covariates. (ii) LDSC estimates are accurate in the absence of covariates, but can become biased in the presence of binary covariates with strong effects. Importantly, regressing covariates out of phenotypes prior to running LDSC can lead to very severe bias, and should always be avoided. We further caution that the software implementation of LDSC can lead to different estimates than those of PCGC-s even in the absence of covariates due to using different data preprocessing procedures, as discussed below. (iii) PCGC accurately estimates all quantities of interest directly or with summary statistics; (iv) standard summary statistics cannot be used to estimate genetic correlation for traits with binary non-genetic risk factors; we propose here a novel formulation of privacy-preserving summary statistics which can be used for this task

When comparing different methods, it is important to distinguish between the underlying mathematics and the software implementation. Even though PCGC-s and LDSC are roughly equivalent in the absence of covariates, the software implementation of PCGC-s is very careful to perform case-control-aware data preprocessing (e.g. avoiding in-sample SNP standardization, and avoid assuming that the diagonal of the kinship matrix is exactly 1.0; Supplementary Material). This can lead to major differences between the estimates of the software implementations in real data analysis. We therefore recommend that researchers use our software implementation of PCGC-s for analysis of case-control studies regardless of the presence of covariates, because it is careful to preprocess case-control data correctly.

An important issue often raised in the context of heritability estimation regards the validity of the assumed model^33,34^. Specific concerns include the difference between “SNP heritability” as measured in studies and “narrow sense heritability” which assumes that all causal SNPs are measured^7,35^, and the potentially larger difference between “narrow sense heritability” under the additivity assumption and the true genetic heritability in the presence of non-additive effects^36^. These concerns are well founded and should certainly be addressed in practice. However, they are not directly related to our study, which focuses on the performance of different methods when the model assumptions (liability threshold model and linear mixed model) hold. We believe our conclusion, that commonly used methods do not give consistently valid results under the standard model assumptions, are of major interest even given the concerns about the validity of the assumptions themselves.

Finally, the LDSC framework includes several techniques not considered in this work: Estimation of the contribution of functional annotations to the liability variance^37^; improved estimation by weighting of summary statistic^16^; and fitting an intercept from the data rather than using a predetermined one^16^; The first technique can be readily adapted into the PCGC-s framework (Supplementary Material). We do not recommend using the other techniques in case-control studies, as the derivations underlying these techniques assume an additive phenotype with genotype-environment independence. Adapting these procedures into case-control studies under a formal theoretical framework remains a potential avenue for future work.

## Supplemental Data

Supplemental Data include 15 figures, 5 tables and mathematical derivations, and can be found online.

## Acknowledgements

This work was supported by the Israeli Science Foundation grants 1487/12 and 1804/16. This study makes use of data generated by the Wellcome Trust Case Control Consortium. A full list of the investigators who contributed to the generation of the data is available from www.wtccc.org.uk. Funding for the project was provided by the Wellcome Trust under awards 076113, 085475 and 090355. We wish to thank the Swedish Bipolar Collection (SWEBIC, PI Mikael Landén) for making data available. The authors thank Noah Zaitlen, Joel Mefford and Na Cai for useful discussions. This collaboration started at the Computational Genomics Summer Institute funded by NIH grant GM112625.

## Web Resources

PCGC-s,https://github.com/omerwe/PCGCs.

